# Inspiratory muscle training for enhancing repeated-sprint ability: A pilot study

**DOI:** 10.1101/2021.01.22.427755

**Authors:** Ramón F. Rodriguez, Robert J. Aughey, François Billaut, Nathan E. Townsend

**Author notes:** Corresponding Author (RFR).

## Abstract

This pilot study examined the effect of inspiratory muscle training (IMT) on repeated-sprint ability and vastus lateralis reoxygenation. Ten recreationally trained subjects were randomly divided into two groups to complete 4 weeks of IMT or Sham (placebo) training. Pre- and post-intervention, a repeated-sprint ability (RSA) test was performed in both normoxia and hypoxia (F_i_O_2_ ≈ 14.5%). Vastus lateralis reoxygenation (VL_reoxy_), defined as peak to minimum amplitude deoxyhaemoglobin for each sprint/recovery cycle, was assessed during all trials using near-infrared spectroscopy. For total work performed, power analysis revealed that for small, medium and large effects (Cohen’s *f*), sample sizes of n = 8, 16 and 90 respectively, are required to achieve a power of 80% at an α level of 0.05. Maximal inspiratory mouth pressure increased in IMT by 36.5%, 95% CI [20.9, 61.6] and by 2.7%, 95% CI [−4.46, 8.8] in Sham. No clear difference in the change of work completed during the sprints between groups were observed in normoxia (Sham −0.805 kJ, 95% CI [−3.92, 0.39]; IMT −2.06 kJ, 95% CI [−11.5, 4.96]; *P* = 0.802), or hypoxia (Sham −3.09 kJ, 95% CI [−7, 0.396]; IMT 0.354 kJ, 95% CI [−1.49, 2.1]; *P* = 0.802). VL_reoxy_ in IMT increased by 9.34%, 95% CI [5.15, 13.7] in normoxia only. In conclusion, despite a large increase in IMT, this was only associated with a small effect on RSA in our pilot study cohort. Owing to a potentially relevant impact of training the inspiratory musculature, future studies should include a sample size of at least 16-20 to detect moderate to large effects on RSA.

## Introduction

During whole-body moderate-intensity exercise, the oxygen cost of breathing contributes 3-6% towards total pulmonary oxygen uptake (VO_2_), which increases to 10-15% during high-intensity exercise [1]. Moreover, if a high work of breathing is sustained, respiratory muscle fatigue can develop, resulting in a reflex increase in muscle sympathetic nerve activity [2]. This response, known as the respiratory muscle metaboreflex, attenuates locomotor muscle blood flow in favour of the respiratory musculature, which hastens the development of locomotor skeletal muscle fatigue [3]. Moreover, exercise in hypoxia is associated with a higher ventilatory equivalent for oxygen and peripheral muscle fatigue [4]. Therefore, respiratory muscle training may represent an effective strategy to alleviate the detrimental effect of sustained high work of breathing during intense exercise, particularly under hypoxic conditions.

Inspiratory muscle training (IMT) has been associated with enhanced exercise performance during the Yo-Yo intermittent recovery test [5, 6], time-trials [7-9], constant load cycling [10, 11], and repeated-sprint exercise (RSE) [12]. By improving the functional capacity of the respiratory muscles, the relative intensity of breathing at a given ventilatory rate decreases. Reducing the relative intensity of hyperpnoea following IMT has been shown to blunt the respiratory muscle metaboreflex [13, 14], reduce the O_2_ cost of breathing [15], and lessen respiratory muscle fatigue in both normoxia and hypoxia [16]. The application of IMT as a method to enhance repeated-sprint ability (RSA) has only been tested in field-based protocols [12, 17], with no work to our knowledge in a controlled laboratory setting under hypoxic conditions.

The ability to maintain performance during RSE is underpinned by the capacity to deliver O_2_ to the locomotor muscles in the short rest periods between sprints [18]. Thus, when RSE is performed in hypoxia, this capacity is negatively impacted [19]. However, respiratory muscle oxygenation appears to be protected, potentially reflecting preferential blood flow redistribution to the respiratory muscles [20]. Interestingly, there is some indication that heightened respiratory muscle work has little consequence on locomotor muscle oxygenation during RSE performed in normoxia [21]. It is possible that when high-intensity exercise is interspersed with rest periods, there is enough capacity in the cardiovascular system to maintain O_2_ supply to both the locomotor and respiratory muscle. Nevertheless, IMT has been shown to improve RSE in normoxic conditions [12], and thus, could also be beneficial for RSE in hypoxia where a higher work of breathing is typically incurred [4]. By enhancing the capacity of the respiratory musculature, the activation of the respiratory muscle metaboreflex may be delayed, thereby improving RSE [15]. We examined data from a previous investigation [20], and carried out a pilot study to determine the feasibility of IMT to induce an ergogenic effect on RSE performance in normoxia and hypoxia.

## Methods

### Power and sample size estimation

Power and sample size estimations were conducted using G*Power 3.1.9.6 [22]. Total work performed on the cycle ergometer was considered the primary outcome measure and was the focus of analysis. Calculations were based on a two-way analysis of variance (ANOVA) with two levels for group (control vs experimental), and two levels for time (Pre-vs Post-intervention). Nonsphericity correction ε was set to 1, and the Pearson’s moment correlation coefficient (*r*) was determined from the second familiarisation and the normoxia pre-intervention trials (Fig 1).

**Fig 1:**
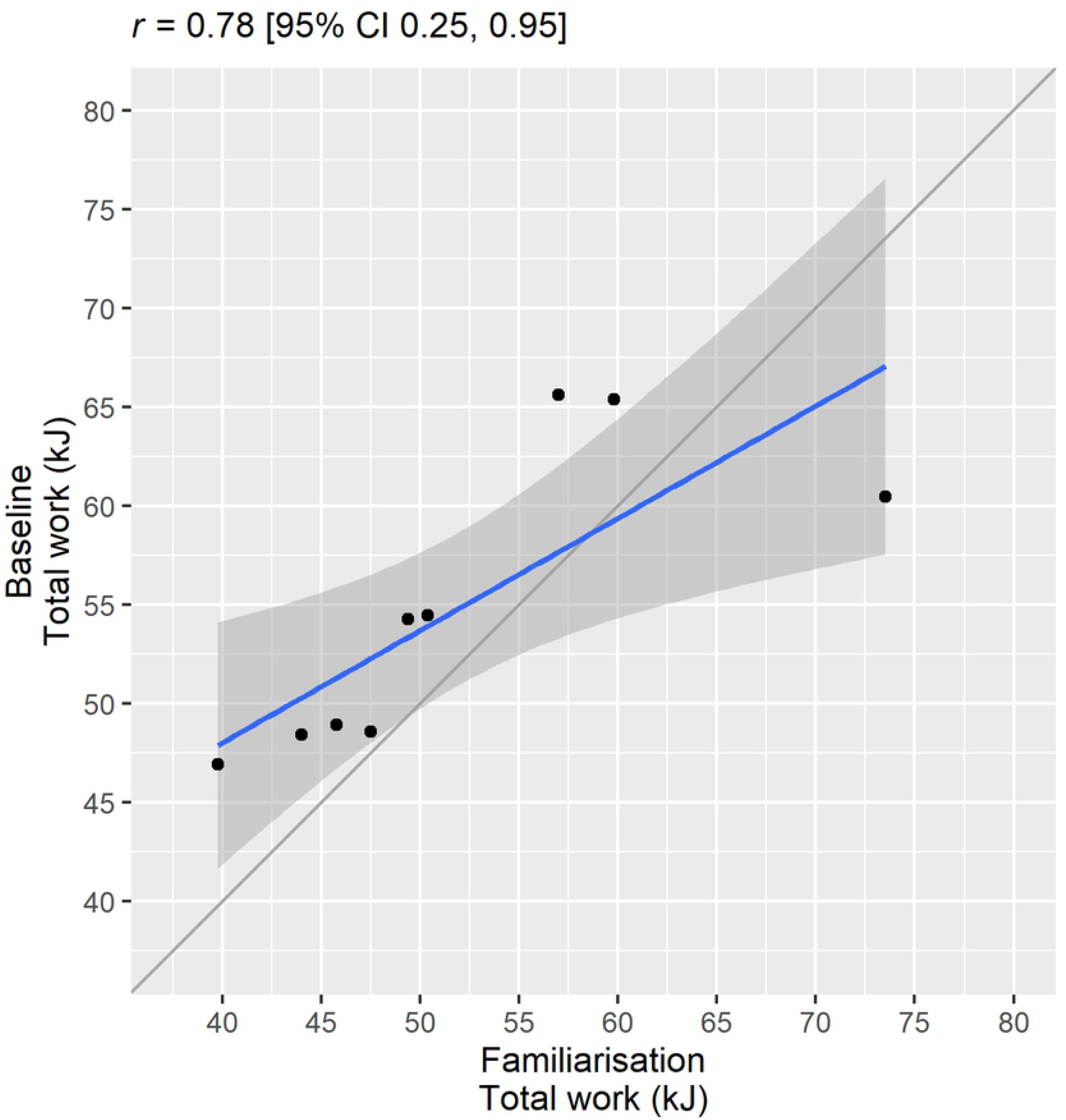
Correlation between the total mechanical work completed during the second Familiarisation trial and the Baseline Normoxia trial. The black dots represent individual subjects (n = 9), and the grey line represents the line of identity. A linear model was fit to the data represented by the blue line and 95% CI by the grey shaded area. Pearson’s moment correlation coefficient (*r*) was calculated and used in the sample size and power analysis as the “correlation among repeated measures”.

Power (1 - β) was calculated as a function of sample size (n) and effect size (Cohen’s *f*). Effect size thresholds were set at small, 0.01; medium, 0.25; large, 0.4. The effect of α error probability was also assessed using at the 0.01,0.05 and 0.1 level.

### Pilot study

#### Design

Ten males accustomed to high-intensity activity were recruited to participate in this study (Sham: age = 24.8 ± 2.4 years, body mass = 77.0 ± 10.3 kg, height = 77.6 ± 6.8 m; IMT: age = 27.2 ± 2.2 years, body mass = 80.2 ± 9.3 kg, height = 179.0 ± 9.0 m). Subjects self-reported to be healthy and with no known neurological, cardiovascular or respiratory diseases. After being fully informed of the requirements, benefits, and risks associated with participation, each subject gave written informed consent. Ethical approval for the study was obtained from the institutional Human Research Ethics Committee, and the study conformed to the declaration of Helsinki.

Participants reported to the laboratory for RSE testing on four separate occasions, which included one session in normoxic (20.78 ± 0.17% O_2_) and normobaric hypoxia (14.49 ± 0.33% O_2_) conditions, both pre- and post-intervention (Altitude Training Systems, Pulford Air and Gas Pty Ltd, Australia). All exercise trials were single-blinded and performed in a counterbalanced order. Testing was conducted within a 23.92 m^2^ environmental chamber set to 21°C and 40% relative humidity (Heuch Pty Ltd, Australia). The training intervention commenced the following day after pre-testing. Post-testing began two days following the intervention period and was completed within 48-72 hours.

#### Inspiratory muscle training

Subjects were randomly assigned to 4 weeks of either Inspiratory Muscle Training (IMT) or Sham training using a POWERbreathe® pressure threshold device (POWERbreathe®, HaB International Ltd, UK). Subjects were naïve that a Sham training group existed, but were informed that the study was investigating the effects of strength (IMT) vs. endurance (Sham) respiratory muscle training. The IMT group completed 30 inspiratory efforts at a pressure threshold starting at 50% of maximal inspiratory (mouth) pressure (MIP), twice per day (AM and PM), every day for 4 weeks. This protocol has been shown to elicit significant improvements in MIP [9, 10, 12, 15]. Once participants could complete 30 breaths comfortably, they were instructed to increase the pressure threshold. The Sham group completed one session per day of 60 breaths at a pressure threshold corresponding to 15% MIP, every day for 4 weeks. The pressure threshold remained at 15% MIP for the entire intervention period, which has been shown to elicit no significant change in MIP [9, 12]. Subjects visited the laboratory weekly for training monitoring and inspiratory muscle strength assessment. A handheld respiratory pressure meter was used (MicroRPM, Micro Medical, Hoechberg, Germany) to measure MIP [23]. Pre-intervention, MIP was assessed to be 133 ± 25 cmH_2_O and 116 ± 41 cmH_2_O in the Sham and IMT groups respectively.

#### Repeated-sprint exercise

Testing was performed on an electromagnetically-braked cycle ergometer (Excalibur, Lode, Groningen, The Netherlands), in isokinetic mode (120 RPM). Subjects completed a 7 min warm-up consisting of 5 min of unloaded cycling at 60-70 RPM and two 4 s maximal effort sprints separated by 1 min each, then rested for another 2.5 min before commencing the repeated-sprint protocol. The RSE protocol included ten maximal sprint efforts lasting 10 s each, separated by 30 passive rest [20].

Therefore, each sprint/recovery duty cycle was 40 s in total. Before the experimental sessions, participants completed two familiarisation trials within one week of commencing the study.

#### Near-infrared spectroscopy

Locomotor muscle oxygenation was measured using NIRS (Oxymon MKIII, Artinis, The Netherlands). The optical sensor was fixed over the distal part of the vastus lateralis muscle belly approximately 15 cm above the proximal border of the patella. Source-detector optode spacing was set to 4.5 cm, and a differential pathlength factor of 4.95 was used [20]. Data were acquired at 10 Hz. A 10^th^ order zero-lag low-pass Butterworth filter was applied to the data to remove movement artefact and signal oscillation due to pedalling [24]. The filtered signal was used for all data analysis thereafter. Vastus lateralis deoxyhaemoglobin (HHb_VL_) was normalised to femoral artery occlusion so that 0% represented a 5 s average immediately prior the occlusion and 100% represented the maximum 5 s average. Arterial occlusion was achieved by placing a cuff around the root of the thigh, which was inflated to 300-350 mmHg until HHb_VL_ plateaued (3-7 min). Peaks and nadirs were identified for each 40 s sprint recovery period, and VL_reoxy_ was calculated as the difference between the peak to nadir of the HHb_VL_ signal.

### Statistical analysis

All data were analysed in the R environment using the estimation statistics framework with the *dabest* package [25], and in addition, repeated measures ANOVA’s were performed on the mechanical work data with the *stats* [26] and *sjstats* [27] packages. Maximal inspiratory pressure (MIP), total mechanical work and average VL_reoxy_ are presented as raw data and effect size (mean difference) with bootstrap 95% confidence interval (95% CI) statistics in Cumming estimation plots. Total work completed and VL_reoxy_ for each 40 s duty cycle, are presented as raw data with mean ± standard deviation.

## Results

### Power and sample size estimation

Adopting the conventionally accepted power of 80% and a 5% α error probability (Fig 2 B), the estimated total sample size to detect a small (0.01), medium (0.25), and large (0.4) effect size for total work is 90, 16 and 8, respectively.

**Fig 2:**
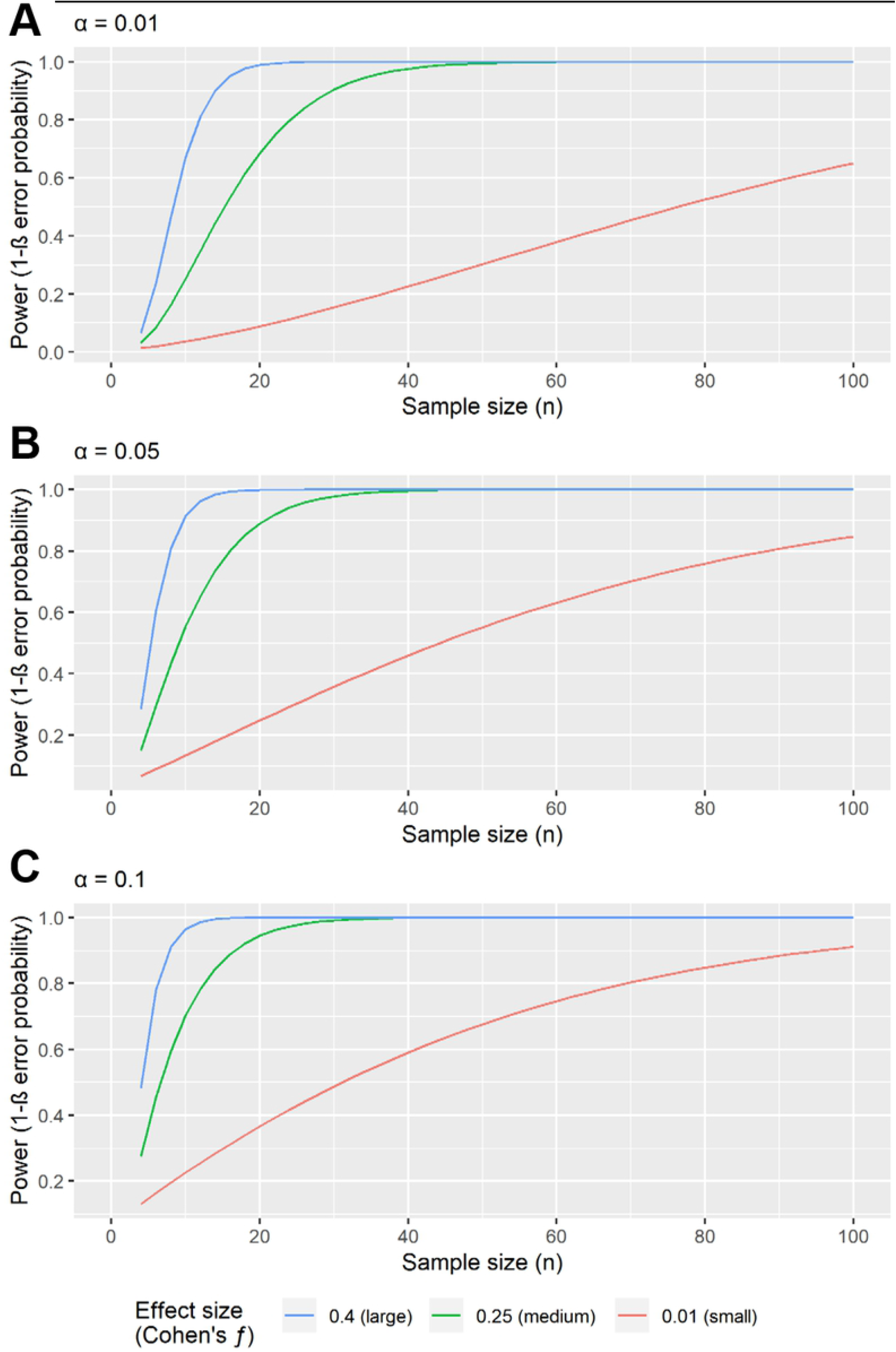
Power as a function of sample size and effect size. Sample size was calculated based on performing an ANOVA: repeated-measures, within-between interaction effects. Nonsphericity correction ε was set to 1, and the correlation among repeated measures was set to 0.78, the number of groups was 2 and number of measurements was 2. Calculations were based on α levels of 0.01 (panel A), 0.05 (panel B), and 0.1 (panel C)

### Pilot Study

After four weeks of training, all participants in the IMP group increased their MIP, which represented a 36.5%, 95% [CI 20.9; 61.6] increase, whereas it remains mostly constant on Sham (2.7%, 95% CI [−4.46, 8.8]). Group mean differences and individual changes are changes are presented in Fig 3.

**Fig 3:**
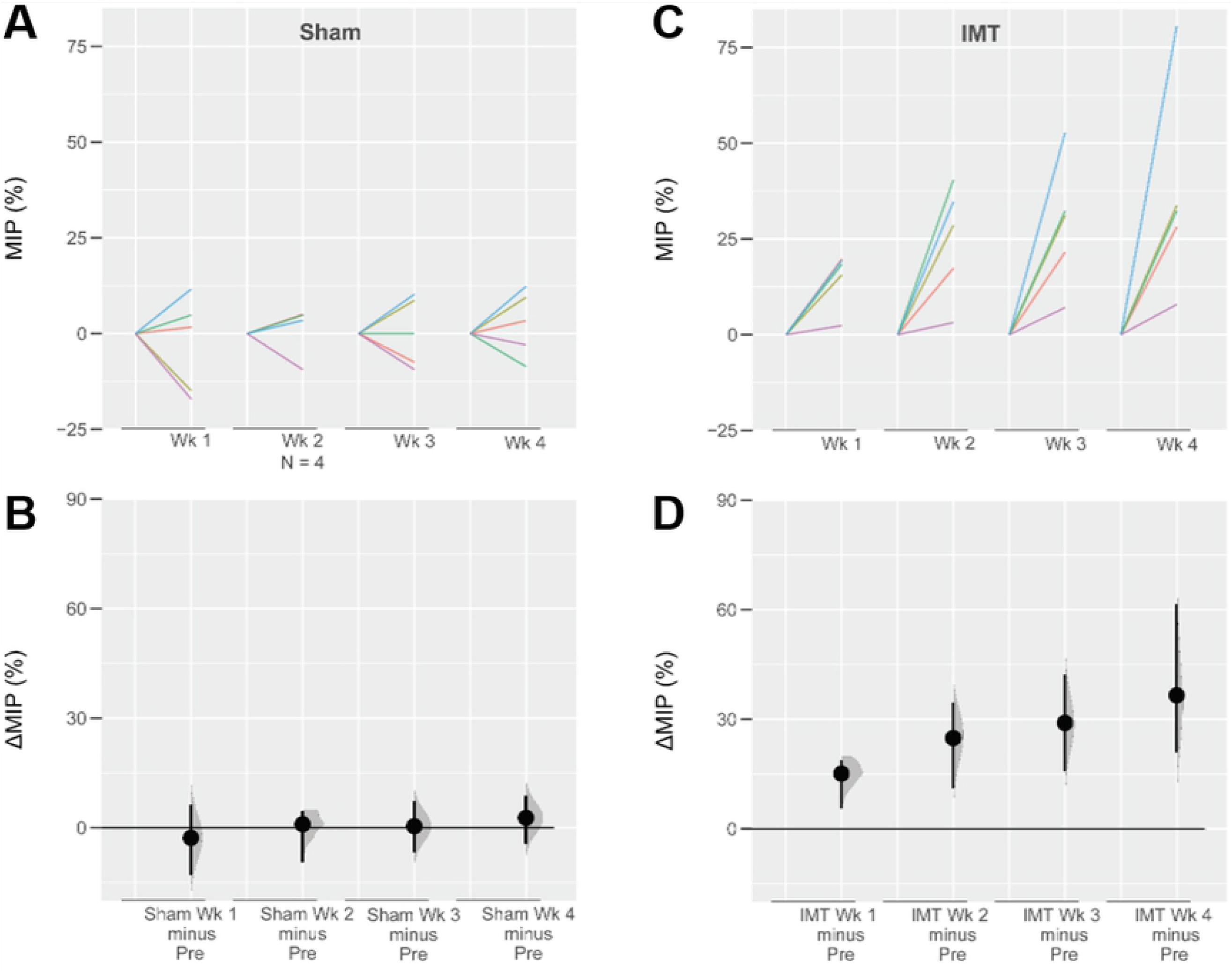
Paired mean difference of the relative change from pre-intervention for maximal inspiratory pressure over the four weeks training period are shown in the above Cumming estimation plot. The raw data is plotted on the upper axes (panels A and C); each mean difference is plotted on the lower axes as a bootstrap sampling distribution (panels B and D). Mean differences are depicted as dots; 95% confidence intervals are indicated by the ends of the vertical error bars.

Outcomes from the ANOVA’s are presented in Table 1 and Table 2. Total work completed and VL_reoxy_ for each trial is presented in Fig 4. There were no clear changes in total work for either normoxia (Sham −0.805 kJ, 95% CI [−3.92, 0.39]; IMT −2.06 kJ, 95% CI [−11.5, 4.96]), or hypoxia (Sham −3.09 kJ, 95% CI [−7, 0.396]; IMT 0.354 kJ, 95% CI [−1.49, 2.1]).

**Fig 4:**
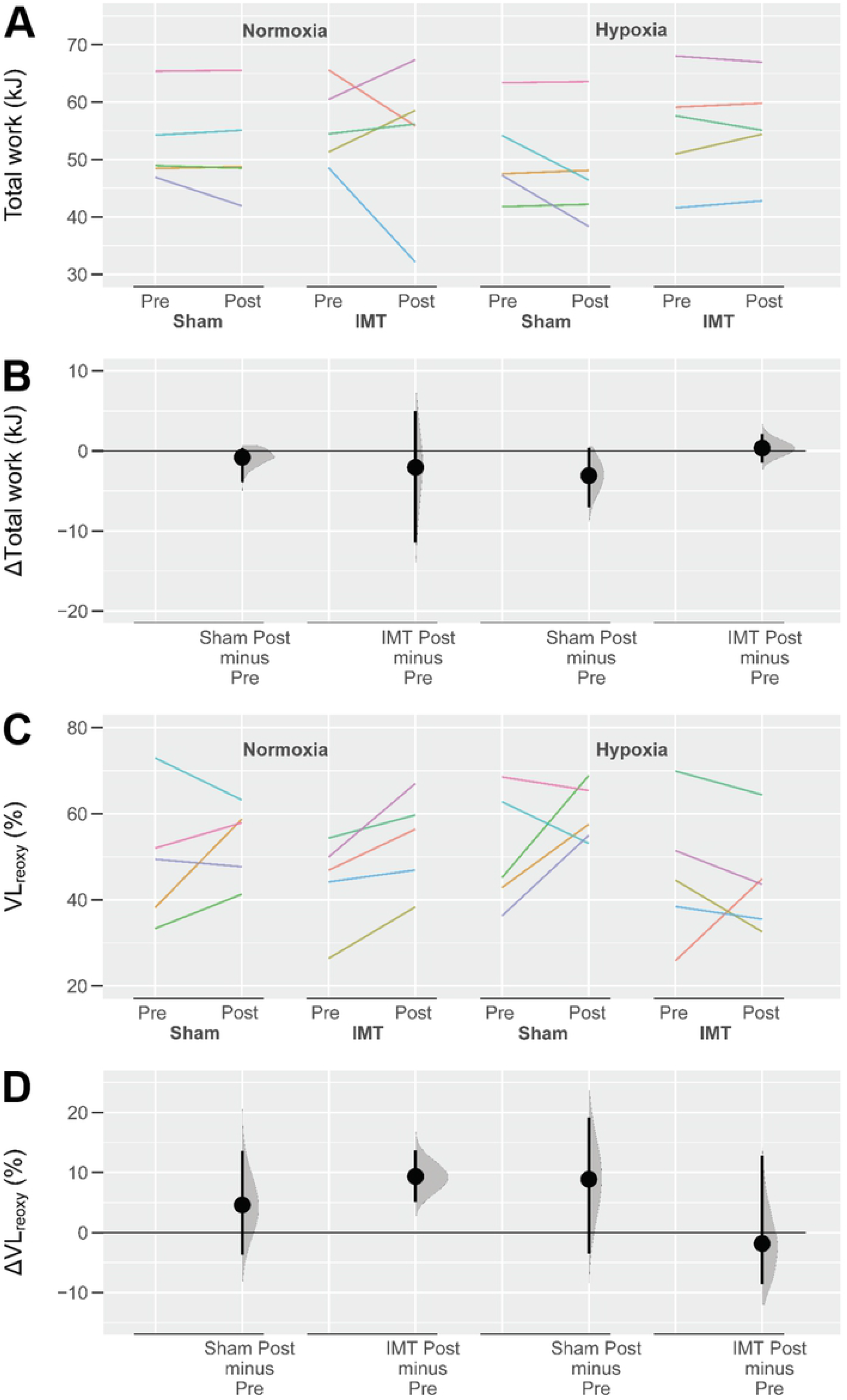
Paired mean difference for 4 comparisons of total mechanical work and VL_reoxy_ is shown in the above Cumming estimation plot. Raw total work data is plotted in panel A, and the mean difference in panel B. Raw VL_reoxy_ is plotted in panel C, and the mean difference in panel D. Mean differences are depicted as dots; 95% confidence intervals are indicated by the ends of the vertical error bars.

**Table 1:** Analysis of variance results for total work performed in normoxia.

| | df | SS | MS | F value | P value | $\eta_p^2$ | $\omega_p^2$ | Cohen's $f$ | Power |
| --- | --- | --- | --- | --- | --- | --- | --- | --- | --- |
| Training (IMT vs. Sham) | 1 | 35.71 | 35.71 | 0.241 | 0.637 | 0.132 | 0.011 | 0.39 | 0.193 |
| Time (Pre vs. Post) | 1 | 10.27 | 10.27 | 0.351 | 0.57 | 0.042 | -0.034 | 0.209 | 0.09 |
| Interaction (Training vs. Time) | 1 | 1.97 | 1.97 | 0.067 | 0.802 | 0.008 | -0.049 | 0.092 | 0.058 |
| Residuals | 8 | 234.319 | 29.29 |  |  |  |  |  |  |

Abbreviations: Degrees of freedom, df; Sum of squares, SS; Mean sum of squares, MS; Partial eta-squared, $\eta_p^2$ ; partial omega-squared, $\omega_p^2$ .

**Table 2:** Analysis of variance (ANOVA) results for total work (kJ) performed in hypoxia.

| | df | SS | MS | F Value | P Value | $\eta_p^2$ | $\omega_p^2$ | Cohen's *f* | Power |
| --- | --- | --- | --- | --- | --- | --- | --- | --- | --- |
| Training (IMT vs. Sham) | 1 | 202.82 | 202.82 | 1.26 | 0.294 | 0.782 | 0.124 | 0.581 | 1.895 |
| Time (Pre vs. Post) | 1 | 9.37 | 9.37 | 1.327 | 0.283 | 0.142 | 0.001 | 0.016 | 0.407 |
| Interaction (Training vs. Time) | 1 | 14.84 | 14.84 | 2.102 | 0.185 | 0.208 | 0.005 | 0.052 | 0.513 |
| Residuals | 8 | 56.50 | 7.06 |  |  |  |  |  |  |

Abbreviations: Degrees of freedom, df; Sum of squares, SS; Mean sum of squares, MS; Partial eta-squared, $\eta_p^2$ ; partial omega-squared, $\omega_p^2$ .

## Discussion

The primary objective of this pilot study was to assess the feasibility of using IMT as a tool for enhancing RSA and improving locomotor muscle tissue oxygenation. Based on the power analysis and sample size estimation of total work, we estimated a sample size between 8 to 90 participants would be required to detect large and small effects, respectively. Considering our sample size of n = 10 (two groups of 5), we should have been able to detect a change in total work if the true effect size was at least large, well beyond the effect sizes (Choen’s *f*appears to be the threshold for exercise intensity at which) observed in the present study of 0.092 and 0.052 Table 1 and Table 2). Given that trained individuals are already well adapted to the demands of high-intensity exercise, it may be that IMT only yields small to moderate effects on performance. Recruiting 90 participants for a training study is not feasible for many exercise science research programs; thus we suggest a total sample size of 16 to 20 participants (two groups of 8 to 10) to provide appropriate statistical power if the effects of IMT on RSE performance is at least moderate.

In the present study, we did not observe any clear performance benefit of IMT in either normoxic or hypoxic conditions (Fig 4). Meta-analysis has demonstrated positive performance benefits of respiratory muscle training (IMT, expiratory muscle training, and both methods combined) for constant-load tests, time-trials and intermittent incremental tests (Yo-Yo intermittent recovery test) [28]. Moreover, linear regression models revealed that test duration also has an important mediating effect on the ergogenic impact of respiratory muscle training. It was estimated that for every minute of exercise, respiratory muscle training provides a 0.4% (95% CI [0.1,0.6%]) performance improvement. Considering our RSE protocol lasts 6 min 10 s, the maximal benefit that we can hope to detect would be 2.5% (95% CI [0.62, 3.7%]). However, there was only 1 min 40 s of actual exercise in our RSE protocol (ten 10 s sprints), and therefore we may only expect a performance benefit of 0.6% (95% CI [0.2, 1%]). The true effect size may lie somewhere in the middle. Moreover, 85% of maximal oxygen uptake (VO_2max_) appears to be the threshold for exercise intensity at which diaphragm fatigue develops [29], and therefore activation of the respiratory muscle metaboreflex [3]. In our previous work [21], we demonstrated that VO_2_ fluctuates between 90% and 70% of VO_2max_ during the sprint and recovery phases, respectively. The time spent above 85% VO_2max_ may not have been sufficient for diaphragm fatigue to develop, and the possible ergogenic effect of IMT to manifest. Prehapse utilising a longer protocol, or one with shorter rest periods relative to the sprint, the benefits of IMT may be more obvious.

We observed a 36.5% increase in inspiratory muscle strength after the 4-week training intervention (Fig 3) which was similar in magnitude to previous studies [9, 12, 15]. Studies of IMT lasting beyond 4 weeks appear to show diminishing effectiveness over time. For example, over an 11-week training period, MIP has been demonstrated to increase by 41% in 4 weeks, and by an additional 4% in the remaining 7 weeks [9]. Previous studies that used a 6-week training intervention demonstrated reduced oxygen cost of hyperpnoea [15]. These results demonstrate that there is a point of diminishing returns of IMT beyond 4 weeks and that extending our training intervention (e.g. to 6 weeks) is unlikely to have yielded different results to what we obtained.

We previously reported that respiratory muscle oxygenation is maintained in hypoxia while VL_reoxy_ is compromised during RSE [20]. We therefore hypothesised that respiratory muscle training could be of potential ergogenic benefit for RSE is hypoxia, as it may reduce the oxygen cost of exercise hyperpnoea, and enhance locomotor muscle oxygen delivery [15, 20]. However, there was no apparent difference in VL_reoxy_ between the training groups (Fig 4 D). Therefore, it may be that respiratory muscle work (oxygen utilisation) has little effect on locomotor muscle oxygenation in RSE. Previously we have demonstrated that despite an increased inspiratory muscle force development, intercostal muscle tissue oxygenation (ration of oxyhaemoglobin to total haemoglobin) can be maintained relative to breathing freely during RSE [21]. The intermittent nature of RSE likely protects against any meaningful competition between the locomotor and respiratory muscles for available oxygen supply.

## Conclusion

These pilot data showed that IMT readily increases the strength of the inspiratory muscles; however, no effect on RSA was found. Based on our sample size calculations, we estimate that we only had the sensitivity to detect a large effect at 80% statistical power. Moreover, to detect medium and small effects, at least 16 and 90 subjects would need to be recruited, respectively. Based on the resources of exercise science laboratories, recruiting 90 subjects may not be feasible. Therefore, we recommend a total sample size of 16-20 is recruited for at least moderate effect sizes to be detected, thus minimising the chances of type II error. Lastly, a double baseline should be utilised to establish the smallest worthwhile change for which the magnitude to the training effect can be judged against.

## Acknowledgments

We would like to thank Mario Popovic for his assistance during data collection. We would also like to thank the laboratory technical staff, Samantha Cassar, Jessica Meilak, and Collene Steward. Funding was provided by the Office for Researcher Training, Quality & Integrity (PhD Student Budget) at Victoria University. The funder had no role in study design, data collection and analysis, decision to publish, or preparation of the manuscript.

## References

1. Harms CA, Wetter TJ, McClaran SR, Pegelow DF, Nickele GA, Nelson WB, et al. Effects of respiratory muscle work on cardiac output and its distribution during maximal exercise. Journal of Applied Physiology. 1998;85(2):609–18. Epub 1998/08/04. doi: 10.1152/jappl.1998.85.2.609.

2. St Croix CM, Morgan BJ, Wetter TJ, Dempsey JA. Fatiguing inspiratory muscle work causes reflex sympathetic activation in humans. The Journal of Physiology. 2000;529(2):493–504. doi: 10.1111/j.1469-7793.2000.00493.x.

3. Dempsey JA, Romer L, Rodman J, Miller J, Smith C. Consequences of exercise-induced respiratory muscle work. Respiratory Physiology and Neurobiology. 2006;151(2-3):242–50. Epub 2006/04/18. doi: 10.1016/j.resp.2005.12.015.

4. Amann M, Pegelow DF, Jacques AJ, Dempsey JA. Inspiratory muscle work in acute hypoxia influences locomotor muscle fatigue and exercise performance of healthy humans. American Journal of Physiology - Regulatory, Integrative and Comparative Physiology. 2007;293(5):R2036–45. Epub 2007/08/24. doi: 10.1152/ajpregu.00442.2007.

5. Lomax M, Grant I, Corbett J. Inspiratory muscle warm-up and inspiratory muscle training: separate and combined effects on intermittent running to exhaustion. Journal of Sports Sciences. 2011;29(6):563–9. Epub 2011/02/25. doi: 10.1080/02640414.2010.543911.

6. Nicks CR, Morgan DW, Fuller DK, Caputo JL. The influence of respiratory muscle training upon intermittent exercise performance. International Journal of Sports Medicine. 2009;30(1):16–21. Epub 2008/11/01. doi: 10.1055/s-2008-1038794.

7. Salazar-Martínez E, Gatterer H, Burtscher M, Naranjo Orellana J, Santalla A. Influence of inspiratory muscle training on ventilatory efficiency and cycling performance in normoxia and hypoxia. Frontiers in Physiology. 2017;8(133):133. Epub 2017/03/25. doi: 10.3389/fphys.2017.00133.

8. Romer LM, McConnell AK, Jones DA. Inspiratory muscle fatigue in trained cyclists: effects of inspiratory muscle training. Medicine and Science in Sports and Exercise. 2002;34(5):785–92. Epub 2002/05/02. doi: 10.1097/00005768-200205000-00010.

9. Volianitis S, McConnell AK, Koutedakis Y, McNaughton L, Backx K, Jones DA. Inspiratory muscle training improves rowing performance. Medicine and Science in Sports and Exercise. 2001;33(5):803–9. Epub 2001/04/27. doi: 10.1097/00005768-200105000-00020.

10. Bailey SJ, Romer LM, Kelly J, Wilkerson DP, DiMenna FJ, Jones AM. Inspiratory muscle training enhances pulmonary O2 uptake kinetics and high-intensity exercise tolerance in humans. Journal of Applied Physiology. 2010;109(2):457–68. Epub 2010/05/29. doi: 10.1152/japplphysiol.00077.2010.

11. Mickleborough TD, Nichols T, Lindley MR, Chatham K, Ionescu AA. Inspiratory flow resistive loading improves respiratory muscle function and endurance capacity in recreational runners. Scandinavian Journal of Medicine and Science in Sports. 2010;20(3):458–68. Epub 2009/06/30. doi: 10.1111/j.1600-0838.2009.00951.x.

12. Romer LM, McConnell AK, Jones DA. Effects of inspiratory muscle training upon recovery time during high intensity, repetitive sprint activity. International Journal of Sports Medicine. 2002;23(5):353–60. Epub 2002/08/08. doi: 10.1055/s-2002-33143.

13. Witt JD, Guenette JA, Rupert JL, McKenzie DC, Sheel AW. Inspiratory muscle training attenuates the human respiratory muscle metaboreflex. The Journal of Physiology. 2007;584(3):1019–28. doi: 10.1113/jphysiol.2007.140855.

14. McConnell AK, Lomax M. The influence of inspiratory muscle work history and specific inspiratory muscle training upon human limb muscle fatigue. The Journal of Physiology. 2006;577(Pt 1):445–57. doi: 10.1113/jphysiol.2006.117614.

15. Turner LA, Tecklenburg-Lund SL, Chapman RF, Stager JM, Wilhite DP, Mickleborough TD. Inspiratory muscle training lowers the oxygen cost of voluntary hyperpnea. Journal of Applied Physiology. 2012;112(1):127–34. Epub 2011/10/08. doi: 10.1152/japplphysiol.00954.2011.

16. Downey AE, Chenoweth LM, Townsend DK, Ranum JD, Ferguson CS, Harms CA. Effects of inspiratory muscle training on exercise responses in normoxia and hypoxia. Respiratory Physiology and Neurobiology. 2007;156(2):137–46. Epub 2006/09/26. doi: 10.1016/j.resp.2006.08.006.

17. Archiza B, Andaku DK, Caruso FCR, Bonjorno JC, Jr., Oliveira CR, Ricci PA, et al. Effects of inspiratory muscle training in professional women football players: a randomized sham-controlled trial. Journal of Sports Sciences. 2018;36(7):771–80. Epub 2017/06/18. doi: 10.1080/02640414.2017.1340659.

18. Buchheit M, Ufland P. Effect of endurance training on performance and muscle reoxygenation rate during repeated-sprint running. European Journal of Applied Physiology. 2011;111(2):293–301. Epub 2010/09/28. doi: 10.1007/s00421-010-1654-9.

19. Billaut F, Buchheit M. Repeated-sprint performance and vastus lateralis oxygenation: Effect of limited O2 availability. Scandinavian Journal of Medicine and Science in Sports. 2013;23(3):185–93. Epub 2013/02/01. doi: 10.1111/sms.12052.

20. Rodriguez RF, Townsend NE, Aughey RJ, Billaut F. Respiratory muscle oxygenation is not impacted by hypoxia during repeated-sprint exercise. Respiratory Physiology and Neurobiology. 2019;260:114–21. Epub 2018/11/20. doi: org/10.1016/j.resp.2018.11.006.

21. Rodriguez RF, Townsend NE, Aughey RJ, Billaut F. Muscle oxygenation maintained during repeated-sprints despite inspiratory muscle loading. PLOS ONE. 2019;14(9):e0222487. Epub 2019/09/20. doi: 10.1371/journal.pone.0222487.

22. Faul F, Erdfelder E, Lang A-G, Buchner A. G*Power 3: A flexible statistical power analysis program for the social, behavioral, and biomedical sciences. Behavior Research Methods. 2007;39(2):175–91. Epub 2007/08/19. doi: 10.3758/BF03193146.

23. American Thoracic Society/European Respiratory Society. ATS/ERS Statement on respiratory muscle testing. American Journal of Respiratory and Critical Care Medicine. 2002;166(4):518–624. Epub 2002/08/21. doi: 10.1164/rccm.166.4.518.

24. Rodriguez RF, Townsend NE, Aughey RJ, Billaut F. Influence of averaging method on muscle deoxygenation interpretation during repeated-sprint exercise. Scandinavian Journal of Medicine and Science in Sports. 2018;28(11):2263–71. Epub 2018/06/09. doi: 10.1111/sms.13238.

25. Ho J, Tumkaya T, Aryal S, Choi H, Claridge-Chang A. Moving beyond P values: data analysis with estimation graphics. Nature Methods. 2019;16(7):565–6. Epub 2019/06/21. doi: 10.1038/s41592-019-0470-3.

26. R Core Team. R: A language and environment for statistical computing. 3.6.3 ed. Vienna, Austria: R Foundation for Statistical Computing; 2020.

27. Daniel L. Statistical Functions for Regression Models. 0.17.9 ed: Zenodo; 2020.

28. Illi SK, Held U, Frank I, Spengler CM. Effect of respiratory muscle training on exercise performance in healthy individuals: a systematic review and meta-analysis. Sports Medicine. 2012;42(8):707–24. Epub 2012/07/07. doi: 10.2165/11631670-000000000-00000. PubMed PMID: 22765281.

29. Johnson BD, Babcock MA, Suman OE, Dempsey JA. Exercise-induced diaphragmatic fatigue in healthy humans. The Journal of Physiology. 1993;460(1):385–405. doi: 10.1113/jphysiol.1993.sp019477.

